# Insights on macrosynteny, ‘rebel’ genes, and a new sex-linked region in anurans from comparative genomics and a new chromosome-level genome for the western chorus frog

**DOI:** 10.1101/2024.10.27.620512

**Authors:** Ying Chen, David R. Lougheed, Zhengxin Sun, Jeffrey Ethier, Vance L. Trudeau, Stephen C. Lougheed

## Abstract

Amphibians have unique genome characteristics including slow karyotypic evolution and cytogenetically undifferentiated sex chromosomes. Yet our understanding of amphibian genomes has not kept pace with that of mammals and birds, partially due to scarce genomic resources and challenges associated with large genome sizes and high repetitiveness. We assembled and annotated a chromosome-level genome for the western chorus frog (*Pseudacris triseriata*), a species of conservation concern and importance in evolutionary research. Comparison of our new genome with other chromosome-level frog genomes reveals exceptionally conserved evolution of 13 chromosomal elements and gene orders across over 200 million years of anuran evolution. We uncovered ‘rebel’ Benchmarking Universal Single-Copy Orthologs (BUSCO) genes that have been duplicated in almost all frog species, have been transposed, and showed lineage-specific synteny patterns – possibly relating to key traits such as frog advertisement calls and mitochondrial genome evolution. We also assembled a complete mitochondrial genome and found heteroplasmy of both point polymorphisms and length variation in the tandem repeat arrays in the control region. Double-digest restriction-site associated DNA sequencing analysis indicates that the western chorus frog has an XY sex system and the sex-linked region involved an ∼1Mb indel structural variant.

Overall, our study provides important genomic resources for treefrogs and other anurans, documents highly conserved chromosomal evolution and gene orders in anurans, identifies ‘rebel’ genes that might be important for frog evolution, and reveals a new sex-linked region with indel structural variants in anurans.

## Introduction

Amphibians have unique genomic characteristics, most notably their large genome sizes, reaching 120Gb in some species (Olmo, 1973; Canapa et al., 2016). Their karyotype evolution features progressive decrease in chromosome number with disappearance of micro- and acrocentric chromosomes (Morescalchi, 1977; Morescalchi, 1980; Sessions, 2008). The evolutionary trends towards a lower number of metacentric chromosomes (through chromosomal fusion) occurred independently in all three amphibian orders of frogs and toads, salamanders and caecilians (Morescalchi, 1980; Nussbaum, 1991; Sessions, 2008). This convergent chromosomal evolution, termed karyotypic orthoselection by White (1973), might be imposed by environmental constraints or by internal cell properties (White, 1973). Stebbins (1966) hypothesized that reduction in chromosome number is adaptive, if environments remained constant, by tightening gene linkage groups on the same chromosomal arms. This hypothesis is partially supported by recent genomic data that showed highly conserved macrosynteny in multiple frog lineages over 200 million years of anuran evolution (Bredeson et al., 2024). Synteny, however, can be conserved as gene linkages on the same chromosomal elements without conserving gene orders (Bhutkar et al., 2008; Simakov et al., 2022).

Genes are not randomly distributed along chromosomes for reasons that are not entirely understood (Hurst, Pál & Lercher, 2004; Michalak, 2008). Genes with essential functions are often clustered into co-expressed neighborhoods, potentially driven by selection favoring lower gene expression noise (Batada & Hurst, 2007; Kustatscher, Grabowski & Rappsilber, 2017) or a consequence of neutral coevolution (Sémon & Duret, 2006). The support for neighboring genes often being functionally related is mixed, with some studies supporting this assertion (Al-Shahrour et al., 2010), others finding no evidence (Sémon & Duret, 2006; Xu, Chen & Shen, 2012). From an ecological perspective, selection for local adaptation might underlie gene rearrangement and physical linkage resulting in genomic islands of differentiation, but direct evidence for this hypothesis is scarce (Yeaman, 2013). Genes that might be more relevant in changing environments (e.g., disease resistance) are more mobile and transpose frequently (Freeling et al., 2008; Woodhouse, Tang & Freeling, 2011). Genes with altered neighborhoods are more likely to undergo expression divergence than genes with conserved neighborhoods due to modified transcriptome environment, contributing to phenotypic divergence and speciation (De, Teichmann & Babu, 2009; De & Babu, 2010). Modelling also supports the hypothesis that genome structure and rearrangements are linked to fitness and adaptation (Crombach & Hogeweg, 2007). Non-adaptive factors like genetic drift and movement of repetitive elements also must play a role in genomic architecture (Koonin, 2009). Regardless, we lack a comprehensive understanding of non-random gene distributions in genomes. Thus, identifying and characterizing gene clustering and synteny continues to be important for gaining insights on genome evolution and organismal diversification.

Amphibian sex chromosomes also exhibit distinct evolutionary features and evolutionary trajectories (Schartl, Schmid & Nanda, 2016). XY systems are most common in amphibians although ZW, WO and other systems do occur (Sessions, 2008; Ma & Veltsos, 2021). In contrast to mammals and birds that have heteromorphic sex chromosomes with degenerate Y and W chromosomes, respectively, amphibians commonly have undifferentiated sex chromosomes and even enlarged Y and W chromosomes (Schartl, Schmid & Nanda, 2016; Ma & Veltsos, 2021), reflecting different drivers of sex chromosome evolution. The miniaturization of Y or W, in the canonical model, is an inevitable long-term consequence of suppressed recombination and gene loss from accumulated deleterious mutations (Charlesworth, Charlesworth & Marais, 2005). Nevertheless, rare occurrences of recombination in occasional sex-reversed individuals (e.g. XY females) are sufficient to maintain Y (or W) homomorphy through new Y haplotype replacing the older version via selection – dubbed the fountain of youth model (Perrin, 2009), observed in species within frog families Hylidae, Bufonidae and Ranidae (Stöck et al., 2013; Dufresnes et al., 2015; Rodrigues et al., 2018; Ping et al., 2022). Another mechanism maintaining sex chromosome homomorphy is rapid sex chromosome turnovers, where new autosomes are frequently co-opted as sex chromosomes (the hot potato model, Blaser et al., 2014; Dufresnes et al., 2015; Furman & Evans, 2016; Jeffries et al., 2018; Ping et al., 2022). Finally, the larger size of Y and W chromosome might be caused by accumulation of heterochromatin (Chalopin et al., 2015a) or from fusion with autosomes (Schmid & Steinlein, 2018).

Because most amphibian species have cytogenetically undifferentiated sex chromosomes, it has proven difficult to identify sex chromosomes until recently, when sex-linked genetic markers could be identified and mapped to a high-quality reference genome (Brelsford, Dufresnes & Perrin, 2016; Lambert, Skelly & Ezaz, 2016; Kuhl et al., 2024). For most species that lack reference genomes, researchers have relied on the western clawed frog (*Xenopus tropicalis*) genome to infer and compare homologous sex chromosome (e.g. Brelsford, Dufresnes & Perrin, 2016), as it was the first amphibian genome to be assembled and is continuously improved (Hellsten et al., 2010; Bredeson et al., 2024). This, however, creates challenges for inferring sex chromosomes in taxa that have experienced fission and fusion chromosomal rearrangements. For example, Z/W chromosomes of *Pseudis tocantis* are likely homologous to chromosomes 4 and 10 of *X. tropicalis* (Gatto et al., 2022); Z/W chromosomes of *Hyla sarda* and *H. savignyi* are homologous to parts of chromosomes 4 and 7 of *X. tropicalis* (Dufresnes et al., 2021). In these cases, a comprehensive picture of homologous chromosomal relationships and rearrangement events among amphibian lineages is necessary to provide evolutionary context for studying sex evolution. To date, amphibians have the fewest available reference genomes among terrestrial vertebrates, impeding evolutionary research in this group including sex evolution (Kosch et al., 2024).

Even less is known of the actual molecular mechanisms underlying sex determination in amphibians (Nakamura, 2009; Ma & Veltos, 2021). To our knowledge, only two frog species have sex-determining genes/regions identified, the W chromosome-specific *dm-w* gene in the African clawed frog *Xenopus laevis* (Yoshimoto et al., 2008; Mawaribuchi et al., 2017; Cauret et al., 2023) and the indel SV region with Y-specific non-coding RNA in European green toad *Bufo(tes) viridis* (Kuhl et al., 2024).

Multiple lines of evidence suggest that genetic pathways underlying amphibian sex development might be flexible, termed ‘developmental systems drift’ (DSD, True & Haag, 2001). For example, the *dm-w* gene in *X. laevis* is a *dmrt1* paralog and is absent in the Z chromosome and all males (Cauret et al., 2020). It is, however, absent in some females of *X. pygmaeus and X. clivii*, is present in some males in *X. pygmaeus, X. clivii* and *X. victorianus* and present in all individuals of both sex in *X. itombwensis* (Cauret et al., 2020). This suggests that the *dm-w* gene is not essential for sex determination in many closely related species of *X. laevis,* and that a different mechanism must have evolved (Cauret et al., 2020). Similarly, the gene *dmrt1* is sex-linked in multiple bufonid, hylid and ranid species (Brelsford et al., 2013; Brelsford, Dufresnes & Perrin, 2016; Ma et al., 2016), but not in other frog species (Tamschick et al., 2015). The observation that there are naturally occurring sex-reversed individuals in wild populations implies that environment might also play a role in sexual development (Rodrigues et al., 2018; Lambert et al., 2019), although the underlying molecular mechanisms remain elusive (Flament, 2016). Overall, our understanding of sex mechanisms in amphibians is rather limited and known sex genes are often located within a small genomic region involving structural variants (SVs) in species with homomorphic sex chromosomes (Mawaribuchi et al., 2017; Kuhl et al., 2024).

Anurans (frogs and toads) comprise the largest of the three orders of amphibians, with over 7,700 species (AmphibiaWeb, 2024). Most species have a biphasic life cycle with larvae (tadpoles) living in aquatic environments and metamorphosing into terrestrial or semi-aquatic adults. Anurans are split into two suborders, the basal, paraphyletic clade Archaeobatrachia and the modern frogs (Neobatrachia) clade (Portik, Streicher & Wiens, 2023). Neobatrachia contains >96% anurans species and has two major superfamilies, Hyloidea and Ranoidea (Portik, Streicher & Wiens, 2023). Currently only 35 chromosome-level frog genomes (∼0.4% of described species) are available on NCBI (September 2024), proportionally lower than most other vertebrate classes such as mammals and birds (Kosch et al., 2024).

The western chorus frog is a terrestrial tree frog (Hylidae, Hyloidea) of the eastern United States and southeastern Canada, one of 18 recognized chorus frog species (Ethier et al., 2021). It is diploid with 12 chromosomes (Schmid et al., 2018), with no evidence of sexually dimorphic chromosomes (James Bogart, pers. comm.). It is of conservation concern in Canada and parts of the USA with many populations declining rapidly in its American and Canadian range (COSEWIC, 2008; Corser et al., 2012). In eastern Canada, trilling chorus frog populations are treated as a single species, the western chorus frog (*Pseudacris triseriata*), with two ‘Designatable Units’ (a biodiversity unit for conservation purposes in Canada, roughly equivalent to ‘Evolutionarily Significant Unit’ in U.S.) acknowledged by the Committee on the Status of Endangered Wildlife in Canada (COSEWIC, 2008). One of these (the Great Lakes / St. Lawrence - Canadian Shield population) is classified as ‘threatened’ (COSEWIC, 2008). The two population designations reflect the distribution of two mitochondrial lineages in Canada that belong to two non-sister mitochondrial lineages (*P. triseriata* and *P. maculata*) that diverged about 10 million years ago (Lemmon, Lemmon & Cannatella, 2007; Lougheed et al., 2020). Recent genomic data found these two lineages overlie a single nuclear genome and thus this is an example of extreme mito-nuclear discordance pattern, likely resulting from historical hybridization and introgression (Lougheed et al., 2020). Furthermore, for the monophyletic clade containing three species, *P. triseriata*, *P. kalmi* and *P. feriarum*, the mtDNA tree topology is incongruent with nDNA topology (Lemmon et al., 2007; Banker et al., 2020), implying multiple hybridization events among those species. Genomic data and a high-quality reference genome are needed to both disentangle the complex evolutionary history of this taxon and facilitate its conservation.

In this study, we assemble and annotate the first chromosome-level reference genome and a complete mitochondrial genome for the western chorus frog (*Pseudacris triseriata*). We leverage this newly high-quality genome and additional chromosome-level frog genomes from NCBI database to: (1) investigate homologous chromosomal evolution in anurans; (2) test whether gene orders are conserved across anuran species; (3) identify ‘rebel genes’ that have duplicated and transposed and evaluate whether these genes underlie frog lineage evolution; and (4) identify the sex system and sex-linked region in the western chorus frog.

## Materials and Methods

### Sample collection and data generation

We collected one male western chorus frog (*P. triseriata*) in Claireville Conservation Area in Ontario, Canada (43.753°N, 79.657°W) in 2022 under an Ontario Ministry of Natural Resources Wildlife Scientific Collector’s Authorization Permit #1099938. We euthanized the individual using benzocaine hydrochloride and removed the tongue, heart, liver, kidneys and one eye. Dissection was completed within 10 minutes to minimize RNA degradation. Tissues were immediately placed in RNAlater® for storage. We also removed both hindlegs, excised the skin, and then snap froze both in liquid nitrogen for later Hi-C lab work. All tissues, along with rest of the body, were stored in a liquid nitrogen dewar for transport to Queen’s University and then transferred to a -80°C freezer until processing.

We extracted high molecular weight DNA from the liver, tongue and heart using a Qiagen MagAttract HMW DNA Kit (Cat. No. 67563) following the manufacturer’s protocol. DNA quantity and sample purity were checked using NanoDrop OD 260/280 ratio (Thermo Fisher Scientific). DNA molecular size was assessed on the Genomic DNA ScreenTape (Agilent). The peak size for the DNA samples was >60kb and 88% of DNA fragments were >20kb. A total of 5-7 μg of DNA was sheared using Covaris g-TUBEs to 15-18 kb with fragment sizes verified on an Agilent TapeStation. Sheared DNA was used as input for library preparation using the SMRTbell Prep Kit 3.0. In total three libraries were prepared (library 1 using liver DNA, library 2 using tongue DNA, and library 3 using pooled tongue and heart DNA). They were set up for sequencing on a PacBio Revio system, loading one library per Revio SMRTCell for a movie time of 24 hours. All library preparations and sequencing were done at The Center for Applied Genomics (TCAG, SickKids Hospital, Toronto, Canada). A Hi-C library was prepared using the two hindlegs using Dovetail Genomics Omni-C Kit (Cat #21005), Illumina Library Module (Cat #25004) and the Dual Index Primer Set #1 (Cat #25010) at the Plateforme d’Analyses génomiques of the Institut de Biologie Intégrative et des Systèmes (IBIS, Université Laval, Québec, Canada). The library was sequenced on a shared lane of Illumina NovaSeq 6000 S4 for paired-end 150bp reads at Génome Québec (Montréal, Québec).

We generated transcriptomic data from the liver, kidney and eye of our focal individual, a liver sample of another male chorus frog collected in 2022 at Cecil Graham Park in Ontario (44.310°N, 76.446°W; OMNR Permit #1099938), and testes samples from two additional individuals collected in 2023 from Merrickville-Wolford, Ontario (44°55′N 75°50′W; OMNR Permit #1105089). We homogenized tissue samples with 0.5mm-2mm stainless steel beads added to 1.5mL tubes in a Bullet Blender Storm 24 (BBY24M, Next Advance). We extracted total RNA for the liver, kidney and eye samples with a DNase treatment using a Qiagen RNeasy Mini Kit (Cat. #74104). The testes RNA was isolated using NucleoZOL and NucleoSpin RNA extraction kits (Cat. #740406.50, Takara). The total RNA for each sample was quantified, and the integrity was assessed, using 5K / RNA / Charge Variant Assay LabChip GX and RNA Assay Reagent Kit (Perkin Elmer) at Génome Québec. RNA Integrity Numbers were between 7.5-8.6 indicating high quality. mRNA enrichment and library preparation were then performed on 250ng of total RNA for each sample using the Illumina Stranded mRNA Prep Kit. The libraries were quantified using the KAPA Library Quantification Kits - Complete kit (Universal) (Kapa Biosystems) and average fragment size was determined using a Fragment Analyzer (Agilent) instrument. Sequencing for the liver, kidney and eye samples was performed on a shared lane of NovaSeq 6000 S4 for paired-end 100bp reads at Génome Québec (Montreal, Québec) and for testes samples at LC Sciences (Houston, Texas) at paired-end 150bp.

### Mitochondrial genome assembly

We used HiFi reads to assemble a complete mitochondrial genome in mitoHiFi v3.2 (Uliano-Silva et al., 2023). We visually inspected and manually curated the resulting mitogenome and the coverage (‘HiFi-vs-final_mitogenome.sorted.bam’ from mitoHiFi) in IGV v2.16.0 (Robinson et al., 2011). We found tandem repeats in the control region (see Results). To check the copy number of the repeats, we ran Tandem Repeat Finder v4.09.1 (Benson, 1999) with recommended default parameters for each HiFi read that mapped onto the mitogenome and plotted the copy number distribution in *ggplot2* (Wickham, 2016) in R v4.1.3 (R Core Team, 2022). We also searched the mitogenome against the nuclear assembly (see next section) using blastn v2.15 (Camacho et al., 2009) to examine possible nuclear mitochondrial DNA segments (NUMTs), i.e., mitochondrial-like sequences flanking with nuclear sequences.

### Nuclear genome assembly

Before assembly, we generated a *k-*mer database at *k-*mer size of 21 from HiFi data in Meryl v1.3 (Rhie et al., 2020) and then estimated genome size, repeat content, and heterozygosity in GenomeScope2 (Ranallo-Benavidez et al., 2020). We also assessed Hi-C data quality with FastQC v0.12.1 (Andrews, 2010) and used fastp v0.23.4 (Chen et al., 2018) to remove adapter sequences (-- detect_adapter_for_pe), reads with 40% of base pairs at phred quality <20 (-q 20, -u 40), and reads shorter than 50bp (-l 50). We assembled the contigs using HiFi + Hi-C phasing with default parameters in HiFiasm v0.19.8- r603 (Cheng et al., 2022). Contigs were scaffolded in YaHS v1.2 (Zhou et al., 2022) using Hi-C contact signals generated from Dovetail Genomics Omni-C pipeline (https://omni-c.readthedocs.io/en/latest/index.html). In brief, the Omni-C pipeline uses bwa mem v0.7.17-r1188 (Li, 2013) to map Hi-C reads onto the contigs and pairtools v1.0.2 (Open2C et al., 2023) to record 5’-most unique ligation events from reads with minimum mapping quality of 40, removes PCR duplicates, and finally uses samtools sort v1.18 (Danecek et al., 2021) to generate the final HiC bam file. For both haplotype assemblies, we tested for contamination using BlobToolKit (Challis et al., 2020) and removed mitochondrial contigs by blasting the assembled mitochondrial genome (see above) against each haplotype assembly. For the more complete haplotype assembly, we used treeval (version galaxy-dev, https://github.com/sanger-tol/treeval/tree/galaxy_dev) to generate Hi-C files and associated coverage tracks and did manual curation using PretextView v0.2.5 (https://github.com/wtsi-hpag/PretextView) and the rapid-curation pipeline (https://gitlab.com/wtsi-grit/rapid-curation), following Howe et al. (2021). We assigned the chromosome numbers based on size. We assessed the quality of the curated genome using gfastats v1.3.6 (Formenti et al., 2022), Merqury v1.3 (Rhie et al., 2020) and BUSCO v5.5.0 tetrapoda_odb10 dataset (Manni et al., 2021).

### Genome annotation

We identified transposable elements (TE) and other repeats in the chorus frog genome using the Earl Grey TE annotation pipeline v4.1.1 (Baril et al., 2024) and Dfam 3.7 curated elements (Hubley et al., 2016). The soft-masked genome was annotated using our RNA-seq data and OrthoDB v11 protein data (Vertebrate partition) using the BRAKER3 pipeline v3.0.8 (Gabriel et al., 2024; Hoff et al., 2016; Hoff et al., 2019; Brůna et al., 2021). BRAKER3 incorporates outputs from multiple programs including HISAT2 (Kim et al., 2019), GeneMark-ETP (Brůna et al., 2024), DIAMOND (Buchfink et al., 2015), Spaln (Gotoh, 2008; Iwata & Gotoh, 2012), StringTie2 (Kovaka et al., 2019), GffCompare (Pertea & Pertea, 2020), AUGUSTUS (Stanke et al., 2006; Stanke et al., 2008) and TSEBRA (Gabriel et al., 2021). In the end, we ran the *best_by_compleasm.py* script to retain all BUSCO (Benchmarking Universal Single-Copy Orthologs) genes in the GeneMark and AUGUSTUS predicted gene sets. We evaluated the annotation by running BUSCO protein mode on the longest isoform per gene against the tetrapoda_odb10 dataset (Manni et al., 2021) and compared it with BUSCO scores of 17 frog annotations in NCBI. The longest isoform per gene was obtained using AGAT v1.4.1 (Dainat, 2024). We also ran OMArk v0.3 (Nevers et al., 2024) to examine possible contamination and dubious proteins in addition to doing a completeness check. Finally, we predicted protein functions of the annotated genes using InterProScan v5.67-99.0 (Blum et al., 2021) and eggNOG-mapper v2.1.12 (Cantalapiedra et al., 2021). The sequence searches in eggNOG-mapper were performed using DIAMOND (Buchfink, Reuter & Drost, 2021) against eggnog ortholog database (Huerta-Cepas et al., 2019). We then parsed these results and did final functional annotation using Funannotate v1.8.17 with tetrapoda_odb10 BUSCO database (Palmer & Stajich, 2020).

### Macrosynteny and ‘rebel’ genes

We conducted macrosynteny analysis on our assembly along with 22 other chromosome-level frog assemblies from NCBI (Fig. 1, also see the list in Table S1). We ran BUSCO v5.5.0 tetrapoda_odb10 dataset against each assembly and identified synteny blocks if 3 or more shared single-copy BUSCO genes (including both complete and fragmented genes) were in the same gene order using custom R scripts (https://github.com/YingChen94/Ptriseriata_genome). We used JCVI (Tang et al., 2015) to visualize the synteny and colored the conserved ancestral chromosomal segments modified from Bredeson et al. (2024).

**Figure 1.**
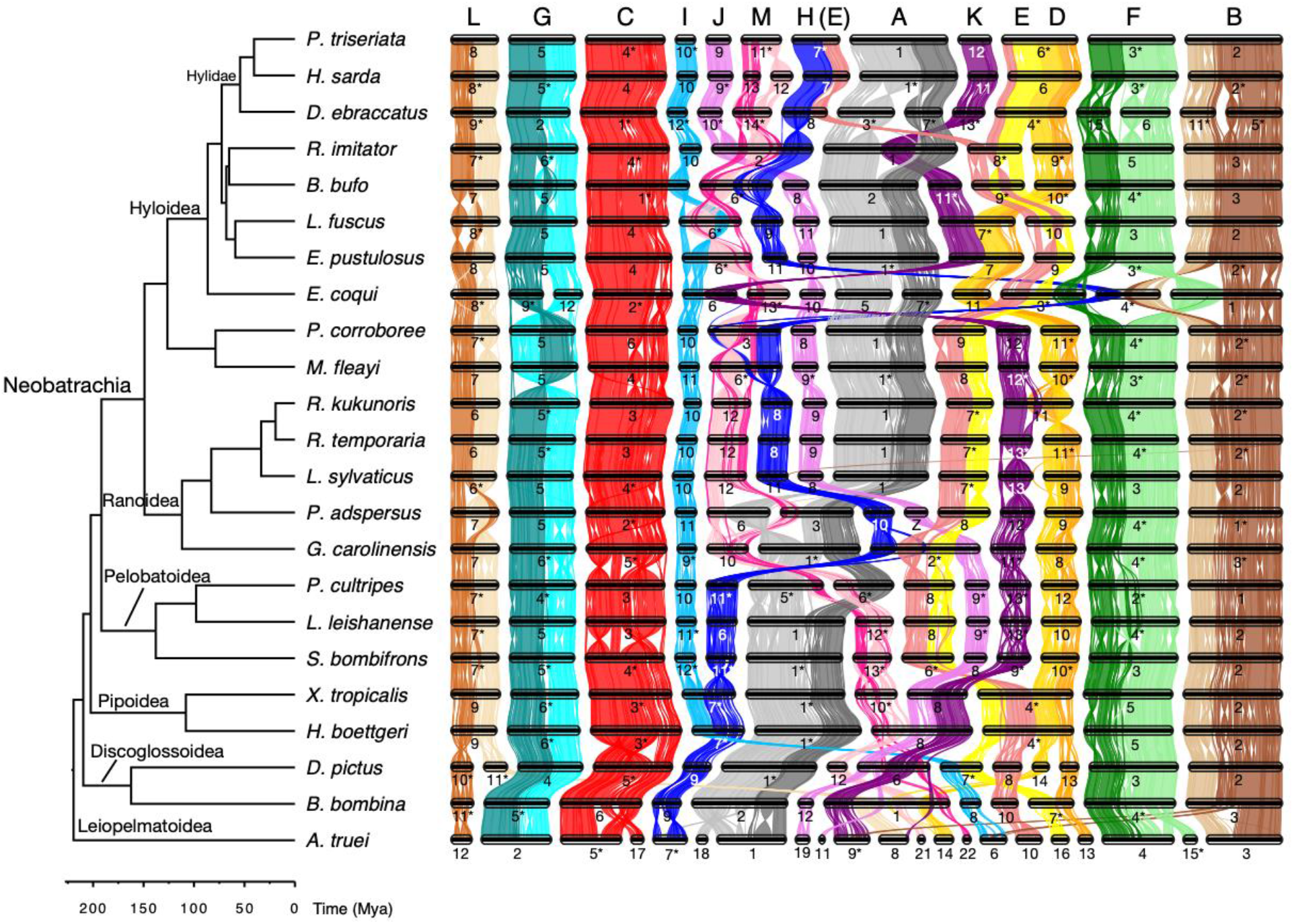
Phylogenetic tree of 23 frog species obtained using timetree (http://www.timetree.org; note that the topology of many Hyloidea families remains unresolved, Portik, Streicher & Wiens, 2023; Streicher et al., 2018; Feng et al., 2017) and a macrosynteny plot, synteny defined as involving three or more BUSCO genes. Chromosomes connoted by asterisks were in the reverse orientation of the original assembly. The length of the chromosome depends on the number of BUSCO genes on them rather than their actual sizes. The 13 conserved chromosomal elements (A-M) are labelled with letters at the top of the syteny plot, following the 13 piparian elements by Bredeson et al. (2024). Note that chromosome 20 of *Ascaphus truei* had too few BUSCO genes to track macrosynteny and thus was not drawn. The accession numbers for each species are: *Ascaphus truei* (GCA_036426205.1, University of California, Berkeley), *Bombina bombina* (GCF_027579735.1, Vertebrate Genomes Project), *Bufo bufo* (GCF_905171765.1, Streicher et al., 2021a), *Dendropsophus ebraccatus* (GCA_027789765.1, Vertebrate Genomes Project), *Discoglossus pictus* (GCA_027410445.1, Vertebrate Genomes Project), *Eleutherodactylus coqui* (GCA_035609145.1, Vertebrate Genomes Project), *Engystomops pustulosus* (GCA_019512145.1, Bredeson et al., 2024), *Gastrophryne carolinensis* (GCA_027917425.1, Vertebrate Genomes Project), *Hyla sarda* (GCF_029499605.1, Vertebrate Genomes Project), *Hymenochirus boettgeri* (GCA_019447015.1, Bredeson et al., 2024), *Leptobrachium leishanense* (GCA_009667805.1, Li et al., 2019), *Leptodactylus fuscus* (GCA_031893025.1, Vertebrate Genomes Project), *Lithobates sylvaticus* (GCA_028564925.1, G10K), *Mixophyes fleayi* (GCA_038048845.1, Vertebrate Genomes Project), *Pelobates cultripes* (GCA_933207985.1, Liedtke et al., 2022), *Pseudophryne corroboree* (GCA_028390025.1, Vertebrate Genomes Project), *Pyxicephalus adspersus* (GCA_032062135.1, Bredeson et al., 2024), *Rana kukunoris* (GCA_029574335.1, Chen et al., 2023), *Rana temporaria* (GCF_905171775.1, Streicher et al., 2021b), *Ranitomeya imitator* (GCA_032444005.1, Vertebrate Genomes Project), *Spea bombifrons* (GCF_027358695.1, Vertebrate Genomes Project), and *Xenopus tropicalis* (GCF_000004195.4, Bredeson et al., 2024).

We further profiled the BUSCO genes with copy number in a phylogenomic context in PanSyn (Yu et al., 2024) and parameters set as k5s5m15 (i.e. 5 top alignment hits for each gene, minimum 5 genes to call a synteny block, and maximum 15 gaps allowed for a synteny block; Zhao & Schranz, 2019). We then conducted network-based microsynteny analysis to test whether any BUSCO genes had lineage specific synteny in the syntenet R package (Zhao & Schranz, 2019, Almeida-Silva et al., 2023). This network method connects syntenic genes into clusters where nodes are genes and edges between nodes represent syntenic relationship (Zhao & Schranz, 2019). Genes that are syntenic across all species will have only one network cluster while transposed genes will not be syntenic and thus will reside in more than one network cluster (Zhao & Schranz, 2019). We identified ‘rebel’ BUSCO genes if they were duplicated in most of frog genomes (since BUSCO genes are expected to be single-copy in most species), or if a gene showed lineage-specific synteny patterns with the criterion that within lineage >90% species were syntenic but between lineages it was not syntenic (i.e., transposed to a different chromosome).

### Sex chromosome identification using ddRADseq data

We performed double digest restriction-site associated DNA sequencing (ddRADseq) on 10 female and 50 male chorus frogs (sample details in Table S2). DNA was standardized to 20 ng/μL and totals of 200 ng per sample were used for ddRAD library preparation at the Plateforme d’Analyses Génomiques (IBIS, Université Laval, Québec, Canada) following Poland et al. (2012) with minor modifications. The restriction enzymes SbfI and MspI were used to digest the DNA with subsequent size selection using a Blue Pippin (SAGE Sciences Inc.) before PCR amplification (elution set between 50 and 65 min on a 2% gel). The library was sequenced on one lane of paired-end 150bp Illumina NovaSeq 6000 S4 at TCAG. We assessed the quality of sequencing in FASTQC v0.12.1 (Andrews, 2010) and demultiplexed the reads using the process_radtags script from the Stacks program v2.64 (Catchen et al., 2013). Reads with uncalled bases were removed (-c) and barcodes and RAD-tags with sequencing errors were rescued (-r).

We used RADSex (Feron et al., 2021) to test whether any ddRAD markers from R1 reads with minimum depth of 10 were associated with sex using Pearson’s chi-squared test of independence, and then mapped those markers to the assembled genome to identify the sex chromosome. We aligned two haplotype assemblies (see above) using minimap2 (Li, 2018) and inspected the sex-linked region in JBrowse2 (Diesh et al., 2023). RNAseq data from six tissue samples (liver, eye, kidney, testes) were mapped onto the reference genome using HISAT2 (Kim et al., 2019) to determine whether any genes were expressed in the sex-linked region.

## Results

### Genome assembly and annotation

The GenomeScope *k*-mer analysis estimated the western chorus frog (*Pseudacris triseriata*) genome to be approximately 3.5Gb in size with 63.5% repetitiveness and a 1.59% heterozygosity rate (Fig. S1A). The final curated haploid assembly was 3.99Gb in size comprising 1,104 scaffolds with a contig N50 of 5.3Mb and scaffold N50 of 441Mb. About 97% of the sequence was assigned to 12 chromosome-level scaffolds (Fig. S1C). BUSCO completeness was 90.4%, among the highest in existing frog genomes (Fig. S2). The per-base consensus accuracy (QV) was 61.7. The *k-*mer completeness was 82.7% for the reference assembly while two haplotype assemblies together reached 98.4% completeness. Results from the Earl Grey pipeline identified 77.3% of the genome as repeats, including 18% classified as DNA transposons, 12.5% as LTR retrotransposons, 6.9% as long interspersed elements (LINEs), 0.2% as short interspersed elements (SINEs), 3.7% as Penelope-like elements and 12.5% as simple repeats, microsatellites and RNA repeats (Fig. S4). A total of 26.9% of the genome was classified as unknown repeats. The BRAKER3 pipeline predicted 22,101 protein-coding genes and 27,426 transcripts, and 57.8% of the genes (n=12,781) were assigned with gene names in the functional annotation in Funannotate. The ratio of mono-exonic to multi-exonic genes was 0.19. The final gene sets had 94.4% BUSCO completeness scores, among the highest of available frog annotations on NCBI (Fig. S5A). The results from OMArk suggested that 88.3% of identified genes were single copy, 3.3% were duplicated and 8.4% had missing hierarchical orthologous groups (Fig. S5B). About 87.1% of proteins were consistent with the tetrapod lineage dataset and no contaminants were reported (Fig. S5B).

### Mitochondrial genome and heteroplasmy

We manually corrected a 27bp mis-assembly in the 16S rRNA region that lacked support from HiFi reads. This assembly error was confirmed by Sanger sequencing the 16S region using the primers 16Sc GTRGGCCTAAAAGCAGCCAC and 16Sd CTCCGGTCTGAACTCAGATCACGTAG (Moriarty & Cannatella, 2004). We found multiple haplotypes in the HiFi reads (Fig. S3A). At position 8302bp in the protein-coding ATP Synthase Membrane Subunit 6 gene (*ATP6*) region, about 56% HiFi reads were ‘G’ and 44% were ‘A’ (Fig. S3A). This is a nonsynonymous substitution predicting either serine (AGC) or glycine (GGC). For the reads with ‘G’ at position 8302bp, almost all reads had ‘C’ at position 15380bp, 15477bp and 15647bp (i.e. G-C-C-C haplotype) except for three HiFi reads having G-T-T-T, G-C-T-C and G-T-T-C. For the reads with ‘A’ at position 8302bp, most reads had ‘T’ at position 15380bp, 15477bp and 15647bp (i.e. A-T-T-T haplotype). Ten reads were A-C-T-T; three reads were A-T-T-C; and the other three reads were A-C-T-C, A-C-C-T, and A-T-C-T. We also found length polymorphisms among the HiFi reads in the two tandem repeat arrays in the control region. The first tandem repeat was 170bp in size and 75.7% HiFi reads supported four copies while 10.1% supported five copies (Fig. S3B). Point mutations were found among HiFi reads in this tandem repeat array (Fig. S3A). The second tandem repeat array (repeat unit = 24bp) had various copy numbers ranging from about 60 to over 100 copies: those HiFi reads with ‘A’ at position 8302bp mostly had a normally distributed copy number centered at 76.5 while the reads with ‘G’ at position 8302bp were more variable (Fig. S3B). We found no evidence of NUMTs and high coverage of HiFi reads supporting two haplotypes (i.e. G-C-C-C haplotype and A-T-T-T haplotype) suggests heteroplasmy. We thus did not curate the mitogenome assembly, which had ‘G’ at position 8302bp (i.e. G-C-C-C haplotype), 4 copies of the 170bp tandem repeat and 99.5 copies of the 24bp repeat. The final mitogenome assembly was 19,745bp with 13 protein-coding genes, 22 transfer RNA genes, two ribosomal RNA genes and one control region. These gene arrangements are typical of Neobatrachia frogs (Zhang et al., 2021).

### Macrosynteny

Macrosynteny was highly conserved across frog species examined, including our focal species, the western chorus frog (Fig. **Error! Reference source not found.**). The most ancient lineage coastal tailed frog (*Ascaphus truei*) had multiple small chromosomes, which have broken and fused into larger chromosomes in *Bombina bombina* and *Discoglossus pictus* and evolved into 13 conserved chromosomal elements. Note that chromosome 6-9 of *A. truei* had small segments fused with larger chromosomes (Fig. 1), which likely involved fission and inversion before fusion. The 13 conserved chromosomal elements went through occasional fusion and fission events but have remained largely intact in many anuran lineages over time without mixing, most likely a result of Robertsonian translocations (Bredeson et al., 2024). Some elements seem to be more likely to fuse: for example, element D and E were fused together twice within Hylidae and Pipoidea, separately; element D and K were fused twice, once in *R. kukunoris* and once in Leptodactylidae; element H and M were fused in Myobatrachidae and also the ancestor of Bufonidae and Dendrobatidae; the fusions of A and M following independent fission events occurred twice, once in *P. cultripes* and the other time in *P. adspersus*. Data from more lineages are needed to test whether such rearrangements occur by chance or are driven by selection. The Hylidae had a unique chromosomal rearrangement where the element E broke into two segments which then fused with D and H (Fig. **Error! Reference source not found.**). Such breakage of a chromosomal element is relatively rare and possibly happened during rapid radiations of families in Hyloidea clade in the late Cretaceous and early Paleocene.

### ‘Rebel’ genes

The phylogenomic profiling plot clearly showed most BUSCO genes were single-copy and syntenic among frog genomes, except for allotetraploid *X. borealis* and *X. laevis* that had many duplicated genes due to polyploidization (Fig. **Error! Reference source not found.**). Two BUSCO genes *nxph2* (neurexophilin 2) and *grsf1* (G-rich RNA sequence binding factor 1) were duplicated in all 29 frog species. Ten BUSCO genes were duplicated in at least 80% of the 29 species: *gabra3* (gamma-aminobutyric acid receptor subunit alpha-3), *fgr* (tyrosine-protein kinase), *aqp4* (aquaporin-4), *ap1s3* (AP-1 complex subunit sigma-3), *slc45a3* (solute carrier family 45 member 3), *acss2* (Acyl-CoA synthetase short chain family member 2), *mcm3* (DNA replication licensing factor MCM3), *chrna9* (neuronal acetylcholine receptor subunit alpha-9), *ctsd* (cathepsin D) and *fam98b* (family with sequence similarity 98 member B). A full list of duplicated or missing BUSCO genes is provided in supplementary Table S3.

About 90.9% of BUSCO genes had a single network cluster, i.e. were syntenic across all frog species with that gene (Fig. 2). About 7.2% BUSCO genes had two network clusters and 1.2% genes had three network clusters (Fig. 2). We identified 34 BUSCO genes that had lineage-specific synteny (supplementary Table S4). Among those, nine genes have two networks split between Archaeobatrachia and Neobatrachia and eight of them were transposed to a different chromosome (supplementary Table S4). Notably, five BUSCO genes that had Archaeobatrachia-specific synteny were in the same genomic region on chromosomal element F of each Archaeobatrachia species: *fgfr1op* (FGFR1 oncogene partner), *b3galnt2* (UDP-GalNAc:beta-1, 3-N-acetylgalactosaminyltransferase 2), *tarbp1* (probable methyltransferase TARBP1), *gpr137b* (integral membrane protein GPR137B), and *chrm3* (muscarinic acetylcholine receptor). The later three of those (*tarbp1*, *gpr137b*, *chrm3*) reside next to each other and were missing in all Neobatrachia frogs, suggesting a deletion event (supplementary Table S4). Seven BUSCO genes had Hyloidea-specific synteny and BUSCO genes had Ranoidea-specific synteny (supplementary Table S4). Notably, gene *twnk* (twinkle mtDNA helicase) and *mrpl43* (mitochondrial ribosomal protein L43) lie adjacent to each other on chromosomal element H in Archaeobatrachia frogs, and then *twnk* transposed to element A in Hyloidea frogs while *mrpl43* transposed to a different chromosome a few times in Neobatrachia frogs (supplementary Table S4).

**Figure 2.**
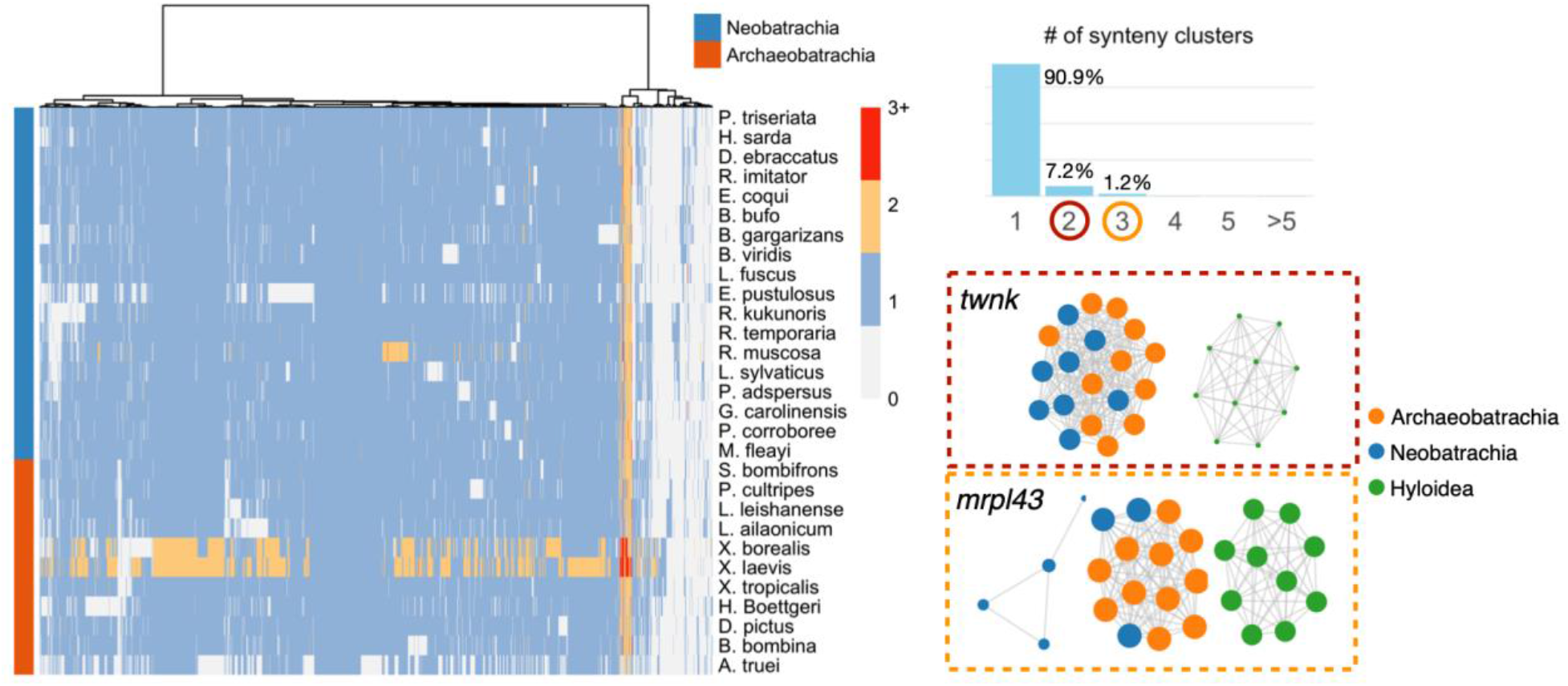
Phylogenomic synteny profiling of 29 frog genomes (left). Most BUSCO genes were single copy and syntenic although many BUSCO genes in *Xenopus borealis* and *X. laevis* had two copies because of polyploidization. The bar plot shows overall percentage of anuran BUSCOs that belong to certain number of synteny clusters (top right). The two synteny networks for gene *twnk* that split between Hyloidea and the rest of species (red dotted box), and three synteny networks for gene *mrpl43* that split between Hyloidea, a few Neobatrachia species and the remaining species (orange dotted box). The accession numbers for some species are provided in the Fig. 1 caption but here with 6 additional species: *Bufo gargarizans* (GCF_014858855.1, Lu et al., 2021), *Bufo(tes) viridis* (GCA_033119425.1, Kuhl et al., 2024), *Leptobrachium ailaonicum* (GCA_032062155.1, University of California, Berkeley), *Rana muscosa* (GCA_029206835.1, Hon et al., 2020), *Xenopus borealis* (GCA_024363595.1, Evans et al., 2022) and *Xenopus laevis* (GCF_017654675.1, International Xenopus Sequencing Consortium).

### Sex chromosomes and sex-linked region

We analyzed a total of 973,056 ddRAD markers present in at least one sample with minimum depth of 10. Among those, 14 markers were significantly associated with males (*p* < 0.05; chi-squared test with Bonferroni correction) and 6 markers were found in all males but no females (Fig. 3A), indicating a male heterogametic sex system. All 14 markers mapped onto the same region between 393,734,341bp to 394,873,959bp on chromosome 1 of the reference genome (Fig. 3B), although 6 additional markers in this region showed no association with sex.

**Figure 3.**
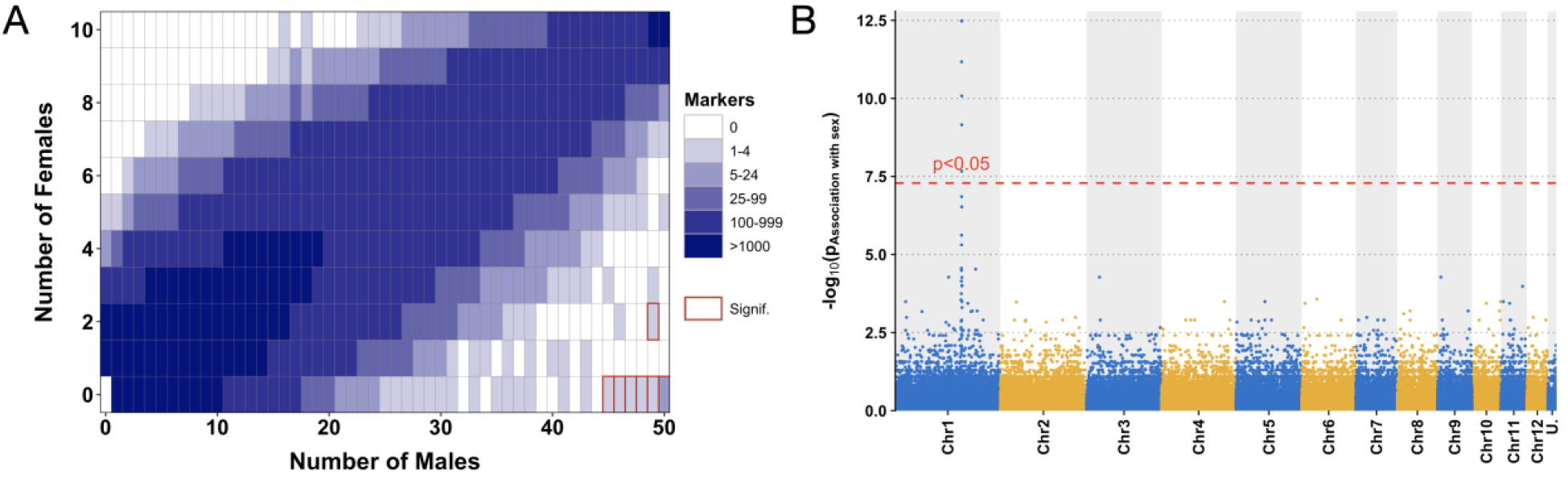
**A.** Tile plots showing the distribution of RADSex markers in male and female chorus frogs. Tiles that are significantly associated with phenotypic males are labelled with red boxes (lower right). **B.** Manhattan plot showing -log10(*P*) of Pearson’s chi-squared test of independence on number of males and number of females, after Bonferroni correction, for all ddRAD markers aligned to the reference genome of the western chorus frog. Unplaced scaffolds were concatenated in a super scaffold (U.) sorted by the order of decreasing size. The dashed line indicates a *p-*value of 0.05 after Bonferroni correction.

Manual inspection of the alignment between two haplotypes revealed structural variation in this sex-linked region (Fig. 4). The Y chromosome (i.e. the haplotype that male-specific ddRAD reads were mapped onto), was larger than the X chromosome (Fig. 4). No gaps were found in this region of the X chromosome suggesting the structural variants in this region were not a result of incomplete assembly. We found the upstream gene *cnot6l* (CCR4-NOT transcription complex subunit 6 like) overlapped with the sex-linked region and was present on both the X and the Y chromosome. *Cnot6l* was expressed in all six RNA tissue samples including two male testis samples. The rest of sex-linked region was mostly composed of repetitive elements including simple repeats, LINEs, hAT transposons, Tc1/mariner transposons, LTRs and unknown repeats.

**Figure 4.**
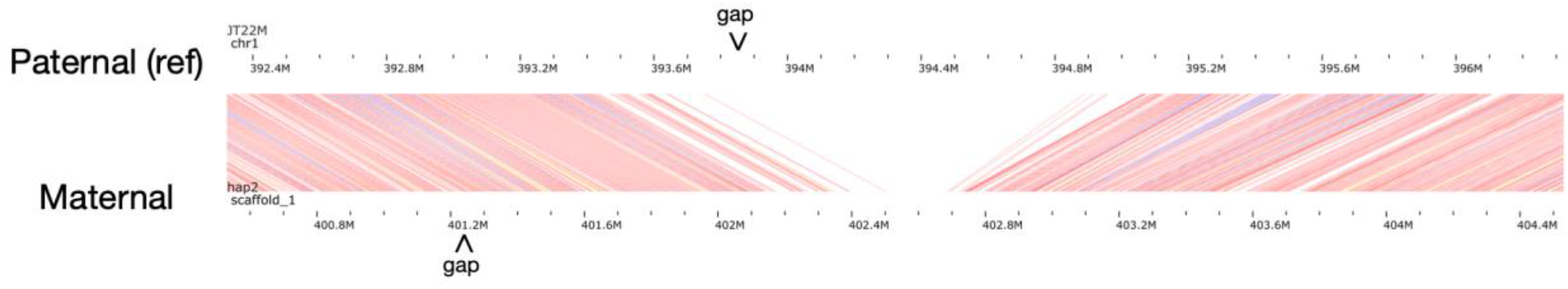
Linear synteny plot revealing lack of synteny in the sex-linked region between paternal Y chromosome (top) and maternal X chromosome (bottom) haplotypes visualized in JBrowse2. Gaps in two assemblies are labelled with arrows.

## Discussion

In this study, we leverage a newly-assembled, chromosome-level chorus frog genome to investigate the evolution of homologous chromosomes in anurans. We found striking macrosyteny and microsyteny across millions of years of anuran evolution. Some individual ‘rebel’ genes may relate to significant diversification and evolution in some clades. We also identified sex chromosomes of chorus frogs and sex-linked region in the western chorus frog.

### Reference genome of western chorus frog

We assembled and annotated a high-quality reference genome for the western chorus frog (*Pseudacris triseriata*), one of only a few genomes in Hylidae, a family comprising over 900 species. More than 75% of the genome consisted of repetitive elements and many repeat activities appear to be recent (Fig. S4). A large proportion of the western chorus frog genome contains DNA transposons and abundant LTRs and LINEs but few SINEs, typical of anuran genomes (Zuo et al., 2023). This contrasts with mammals where there is a predominance of LINEs, SINEs and LTRs but few active DNA transposons with some exceptions like the vespertilionid bats (Pritham & Feschotte, 2007; Chalopin et al., 2015b; Platt II, Vandewege, & Ray, 2018). The evolutionary significance of this is unclear but possibly relates to competition between TEs and host defense mechanisms (Le Rouzic & Capy, 2006; Abrusán & Krambeck, 2006). For the functional components, the chorus frog genome was predicted to encode 22,101 protein-coding genes, comparable to 25,106 genes in *X. tropicalis* and other frog species (Bredeson et al., 2024). The genome of *P. triseriata* can serve as an important genomic resource for research on amphibian evolution and in conservation efforts.

### Heteroplasmy in the mitochondrial genome

We assembled the first complete mitochondrial genome for *P. triseriata*, which included 37 genes in the typical gene order for Neobatrachia frogs (Zhang et al., 2021). We found two tandem repeat arrays in the control region: the first was 170bp in size and, for this individual repeated four times in 75.7% HiFi reads; the other was 24bp in size with highly variable copy number from 60 to over 100 copies (Fig. S3). Some of the copy number variation might come from PacBio insertion and deletion sequencing errors (Wenger et al., 2019), but this cannot explain the large deviation in the second repeat array. We also could not find any evidence of NUMTs and concluded that the repeat variation was true variation within this single individual (i.e., heteroplasmy). Indeed, variable numbers of repeats in the control region are common across vertebrates and invertebrates and many are also heteroplasmic within individuals (Lunt, Whipple & Hyman, 1998). The evolution of the repeat arrays and heteroplasmy can be explained by the ‘illegitimate elongation model’ (Buroker et al. 1990). This model states that the gain and loss of repeat units results from a dynamic competitive base pairing between the L-strand and the pre-existing H-strand or the nascent D-loop strand during mitochondrial DNA replication. If the D-loop strand is partially displaced by the H-strand, the repeat units can form stable hairpin loops, shortening the strand. The subsequent reinvasion of the shortened D-loop strand likely will misalign with an upstream copy of the repeat, resulting in an additional copy in the D-loop strand. Alternatively, if hairpin loops are formed on the H- and L-strand, the D-loop strand misaligning to a downstream copy of the repeat will result in a loss of copy. This model predicts that the central repeats evolve in a concerted manner, and repeat units can diverge among lineages but remain identical within lineages (Buroker et al., 1990). This is supported by the tandem repeat arrays in the control region of a congeneric chorus frog species *P. maculata* where the repeat unit diverged from *P. triseriata* by a few base pairs but remained almost perfectly identical in the array (Chen & Lougheed, unpublished).

Heteroplasmy was also evident in two main haplotypes with point mutations at four positions (see results). The substitution of ‘A’ vs ‘G’ in *ATP6* region was non-synonymous and each encoded amino acids serine (AGC) and glycine (GGC), respectively. By searching in NCBI, we found that most frogs had ‘G’ in this position, while a few species (e.g. *Hyla arenicolor* - NCBI accession HM152156.1; *Osteocephalus planiceps* - NCBI accession OP572177.1) had ‘A’, implying both amino acids were functional. For our annotated genome we cannot know whether this *ATP6* heteroplasmy was inherited or arose *de novo* nor do we know what the ancestral state is in this species; knowledge of the ancestral state can be useful to infer evolutionary processes such as selection (e.g. Li et al., 2016). Nonetheless, heteroplasmy is a necessary first step before novel haplotypes can be fixed and thus these data can be insightful in understanding mitochondrial genome evolution.

Only through manual curation of the mitogenome assembly using alignment of HiFi reads did we identify the 27bp mis-assembly and reveal the extreme intra-individual heteroplasmy. We thus recommend manual curation on mitochondrial assembly which is not only important for accuracy but also to provide insights on heteroplasmy and mitochondrial genome evolution.

### Conserved macrosyteny and gene orders in anurans

The macrosyteny analysis among 23 frog genomes revealed the conserved evolution of 13 ancestral chromosomal elements throughout anuran evolution (Bredeson et al., 2024). We gained insights on how the chromosomes in the most ancient lineage *Ascaphus* was rearranged resulting in decreased chromosome numbers and loss of microchromosomes in more modern lineages (Morescalchi, 1980). Multiple fissions must have happened before fusion events (chromosome 6-9 in *A. truei*), indicating frequent chromosomal rearrangements in these early lineages in contrast to later slow evolution. Strikingly, not only were the same genes retained in each chromosomal element, but gene orders also conserved throughout the evolution with over 90% BUSCO genes being syntenic across all species examined. This is in sharp contrast to the genome evolution of other lineages like the Muller elements in *Drosophila* that had conserved gene linkages on each chromosomal arm but reshuffled gene- orders (Bhutkar et al., 2008). Such gene-order reshuffling is a result of accumulating intra-chromosomal rearrangements like inversions and translocations (Simakov et al., 2022), which can also be observed occasionally in anurans – for example, element K in *Pelobates cultripes* and element M in *Leptobrachium leishanense* (Fig. 1). It is, however, apparent that the conserved gene order in anurans is the dominant pattern. Assuming chromosomal rearrangements are mostly evolutionarily neutral then more diverged lineages should have fewer similarities in gene order (e.g. Bhutkar et al., 2008). The exceptional preservation of both chromosomal elements and gene orders in anurans thus must reflect strong selection. One evolutionary constraint can come from developmental regulatory networks controlling gene expressions (Engström et al., 2007; Irimia et al., 2012). Anurans are known to have “extreme morphological uniformity” and are “the most easily diagnosed of all vertebrate groups” (Griffiths, 1963).

We can speculate then that the conserved chromosomal architecture and gene order underlie the slow body form evolution and biphasic life history in frogs and toads.

### Evolutionarily significant ‘rebel’ BUSCO genes

BUSCO (Benchmarking Universal Single-Copy Orthologs) genes are used as a genome assessment metric because they are conserved as single copies in at least 90% species within tetrapod lineages (Simão et al., 2015). We found some BUSCO genes duplicated in almost all frog species, and we hypothesize that these have played a significant role in anuran trait diversification. For example, anurans are among the most ancient lineages to use acoustic communication (Kelley, 2022) and almost all species have mate recognition systems with advertisement calls except the ancient lineages Ascaphidae and Leiopelmatidae and a few others like *Acanthixalus* and *Megaelosia*. The temporal attributes of their calls (e.g., pulse rate and duration) are involved in sexual selection and premating isolation (Gerhardt, 2005; Lemmon & Lemmon, 2010) and are mediated by the interplay of excitatory inputs via ionotropic glutamate receptors and inhibition via gamma-aminobutyric acid type A (GABAA) receptors in the central nervous system (Alluri et al., 2016; Alluri et al., 2021). GABAergic inhibition influences frequency tuning and temporal selectivity in frog midbrain neurons (Hall, 1999; Alluri, et al., 2021). In the upland chorus frog (*Pseudacris feriarum*), multiple GABA transporter genes are expressed differentially between populations with diverged temporal call attributes of pulse rate due to reinforcement (Lemmon & Lemmon, 2010; Ospina et al., 2021). The ‘rebel’ gene GABAA receptor subunit α3 (*gabra3*) is important in neuronal development (Ohlson et al., 2007; Daniel et al., 2011) and has been implicated in locomotor behavior in zebrafish (Barnaby et al., 2022) and both affective and cognitive functions in mice (Fiorelli et al., 2008). We thus hypothesize that the three to four gene copies of the *gabra3* gene in most frog genomes (except for *Rana muscosa* and *Leptobrachium ailaonicum* with two copies and one copy, respectively) have played a role in neuronal development and calling behaviors in ancestral frogs and are maintained throughout anuran diversification. We found another rebel gene, *aqp4*, to have been duplicated in 27 out of 29 species; it encodes water channel protein aquaporins (AQP) that function in osmoregulation and thermal tolerance and thus might be important in anuran biphasic life histories involving phases in both aquatic and terrestrial environments (Krane & Goldstein, 2007; Suzuki et al., 2007).

We found 34 ‘rebel’ genes with lineage specific synteny patterns and some of them are likely involved in lineage diversification. For example, three ‘rebel’ genes were related to mitochondrial functions: ATP-binding cassette sub-family B member 10 (*abcb10*), twinkle mtDNA helicase (*twnk*) and mitochondrial ribosomal protein L43 gene (*mrpl43*). Gene *abcb10 was* duplicated and transposed in the western spadefoot toad (*Pelobates cultripes,* supplementary Table S4), a species where tadpoles can accelerate their development by ∼32% in response to pond drying at the cost of oxidative stress (Gomez-Mestre, Kulkarni & Buchholz, 2013). The ABCB10 protein can modulate oxidative stress from ischemia/reperfusion (Liesa et al., 2011; Liesa, Qiu & Shirihai, 2012) and the extra copy of *abcb10* might underlie some aspects of the high developmental plasticity in western spadefoot toad. *Twnk* and *mrpl43* lie adjacent to one another in archaeobatrachians; such tight linkage has also been reported in many other vertebrate species like mice and cattle where two genes were arranged head-to-head and shared a bidirectional promoter (Tyynismaa et al., 2004; Meersseman et al., 2014). This gene linkage, however, is lost in neobatrachian frogs (supplementary Table S4), the evolutionary ramifications of which remain to be investigated. We hypothesize that the transposition of *twnk* and *mrpl43* might affect neobatrachian mitochondrial evolution, especially because TWINKLE proteins help to regulate mitochondrial DNA replication (Jemt et al., 2015) and *twnk* and *mrpl43* are associated with mitochondrial heteroplasmy in humans (Nandakumar et al., 2021). Neobatrachians share a synapomorphic feature of a rearranged gene order of LTPF tRNA clusters (*trnL-trnT-trnP-trnF*) in the mitochondrial genome while most archaeobatrachians have the typical vertebrate mitochondrial gene order (Irisarri et al., 2012; Zhang et al., 2021). Other types of mitochondrial gene rearrangements, such as duplications and/or losses of tRNA genes and control regions, are also present in neobatrachians (Su et al., 2007; Kurabayashi et al., 2008) implying a relaxation of purifying selection (Irisarri et al., 2012). It is plausible that disruption of gene linkage between *twnk* and *mrpl43* provided flexibility in mitochondrial evolution.

Functions of many other ‘rebel’ BUSCO genes in anurans are unknown. For example, we found that gene sets *tarbp1*, *gpr137b* and *chrm3* occur next to each other in all surveyed archaeobatrachians but were missing in all neobatrachians. We also found transposition of nearby genes *fgfr1op, b3galnt2* and *sprtn* implying that this genomic region in chromosomal element F was evolutionarily unstable for the ancestor of neobatrachians. Our knowledge of those genes is limited and largely confined to model organisms (Gloriam, Fredriksson & Schiöth, 2007; Pedersen, Bergqvist & Larhammar, 2018) making it difficult to extrapolate their functional roles. Future studies can use this ‘rebel’ gene list as a foundation for studying anurans evolution. We also recognize that rearrangements of some ‘rebel’ genes may be neutral with no significant evolutionary consequences but rather a result of drift or repetitive elements activities (Koonin, 2009). Considering the extreme conservation in gene order in anurans, these ‘rebel’ gene rearrangements, if truly neutral, still warrant investigations for insights in what factors maintain or disrupt the conserved genomic architecture.

### Sex chromosome evolution

Our analysis using ddRADseq data revealed that western chorus frog has XY sex system and that the sex chromosome was the largest in our assembly – chromosome 1 (corresponding to chromosome 1 in *X. tropicalis* and the conserved anuran chromosomal element A; see Fig. **Error! Reference source not found.**). This chromosome pair is thus far the most common sex chromosome pair in anurans (Miura, 2017; Ma & Veltos, 2021), and is present in at least seven hylid species (Dufresnes et al., 2015; Brelsford, Dufresnes & Perrins, 2016; Dufresnes et al., 2021), one pelodryadid species (Bertola et al., 2023), seven species in the family Bufonidae (Tamschick et al., 2015; Kuhl et al., 2024), and two dicroglossid species (Xiao et al., 2024). This has been attributed to the presence of key sex determination genes like *dmrt1* and *Amh* (Brelsford et al., 2013). However, those genes are not the sex-determining master gene for many species (Tamschick et al., 2015) and other chromosomes also harbor candidate sex-determining genes including *Sox3* on chromosome element J and *Cyp17* on element H (Fig. **Error! Reference source not found.**; Miura, 2017). The identity of sex chromosomes is particularly evolutionarily labile in Ranidae and Pipidae, but appears conserved across Hylidae and Bufonidae, except for *H. sarda* and *H. savigni* (Miura, 2017; Dufresnes et al., 2021; Ma & Veltos, 2021). This implies variable evolutionary rates in sex chromosome evolution among anuran lineages. We recognize that the number of anuran species with known sex chromosomes is still too few to make strong assertions, and more data are needed to corroborate these evolutionary patterns.

### Structural variation in sex-linked region

We found a ∼1Mb indel structural variant in the sex-linked region in chromosome 1 where either the putative Y chromosome had an insertion, or the X chromosomes had a deletion. This region was annotated with many repetitive elements and such accumulation of heterochromatin in the sex chromosome can relate to early stages of sex-chromosome differentiation (Schmid et al., 1991; Schartl, Schmid & Nanda, 2016). For example, in Hamilton’s frog (*Leiopelma hamiltoni*), the W chromosome is slightly larger, with more heterochromatin at the centromeric region, relative to the corresponding Z chromosome (Green, 1988). In the edible frog (*Pelophylax esculentus*), the X and Y chromosomes are mostly homomorphic with the Y chromosome differing from the X by a single late replication band (revealed using a bromodeoxyuridine immunochemical assay), likely due to presence of repetitive sequences on the Y chromosome (Schempp & Schmid, 1981). Such accumulation of heterochromatin can reduce recombination rate and prevent meiotic pairing and crossing over (Schartl, Schmid & Nanda, 2016), further facilitating accumulation of repetitive elements and growing structural divergence.

Alternatively, the structural variation in the Y chromosome may play a functional role in sex determination. This has been reported in the European green toad, *Bufo(tes) viridis*, that has an XY system and uses the same sex chromosome pair as the western chorus frog (Kuhl et al., 2024). The Y chromosome of *B. viridis* also contains repetitive elements and a 5.8kb long non-coding RNA (ncRNA) in the 5’ regulatory region of gene *bod1l* (Kuhl et al., 2024). This Y-specific ncRNA was expressed exclusively in males and the expression was highest in testis relative to other organs, suggesting a role in sex-specific differentiation. This Y-specific ncRNA also affected the expression of the gene *bod1l* in males as an enhancer element, evidenced by sexually distinct spatial and temporal expression of *bod1l* in developing gonads of males and females coinciding with Y-specific ncRNA expression (Kuhl et al., 2024). Based on these findings, we hypothesize that the observed structural variation in the sex-linked region of western chorus frogs plays a regulatory role in sex determination. We found that *cnot6l* gene partially overlapped the sex-linked region (note that the exact boundary of sex-linked region cannot be defined due to shorter reads and reduced representation nature of ddRADseq data). *Cnot6l* is a paralog gene of *Cnot6* and they encode protein CNOT6/6L (or CCR4), a catalytic subunit of the Carbon Catabolite Repression—Negative On TATA-less (CCR4-NOT) complex and a key regulator of eukaryotic gene

expression (Collart, 2016). In the model organism, *Mus musculus*, *cnot6l* plays a significant role in sex development by regulating mRNA turnover through deadenylation. For example, *cnot6l* was highly expressed in mouse oocytes compared with other cell types and was critical in meiotic maturation of oocytes (Sha et al., 2018). Deletion of the *Cnot6l* gene (*Cnot6l*^−/−^) led to ∼40% lower fertility in female mice (Horvat et al., 2018) while the double knockout of *Cnot6/6l* genes (*Cnot6/6l*^−/−^) led to female infertility due to defective follicular development (Dai et al., 2021). The functional role of *Cnot6l* in sexual development in anurans is still unknown, but possibly it interacts with the sex-linked region and affects sex development through regulating mRNA expressions in developing gonads of the western chorus frog.

## Conclusion

We assembled and annotated a chromosome-level reference genome and a complete mitochondrial genome for the western chorus frog (*Pseudacris triseriata*). For the mitochondrial genome, heteroplasmy was evident, with two haplotypes comprising point polymorphisms at multiple positions as well as length polymorphisms in the repeat array. Macro- and microsynteny analyses of 29 frog species revealed conserved evolution of both 13 ancestral chromosomal elements and gene orders. We identified some ‘rebel’ BUSCO genes that were duplicated in most frog genomes and had lineage-specific synteny patterns, possibly playing a significant role in anuran trait evolution, mitochondrial function, and diversification. Using ddRADseq data, we found an XY sex system in the western chorus frog, with the sex chromosome pair conserved in Hylidae and Bufonidae lineages. We report a new sex-linked region with indel structural variation between X and Y chromosome, possibly determining sex by interacting with the upstream gene *cnot6l*. Our work provided novel genomic resources for an anuran species and family of conservation concern and evolutionary interest, documented the homologous chromosome relationships among 23 frog genomes and their extremely conserved gene orders, compiled a list of ‘rebel’ genes that warrants future research, and revealed a new sex-linked region that contributing to our understanding of anuran chromosomal and sex evolution.

## Acknowledgements

This work was supported by Environment and Climate Change Canada and the Natural Sciences and Engineering Research Council of Canada (NSERC) via Discovery Grants to S.C.L. and V.L.T. We are grateful to John Brett, Jennifer Thompson and Madeleine Robitaille for their help with fieldwork and Stacey Robinson for her help on RNAseq experiments. We thank the Toronto and Region Conservation Authority for the sampling permission at the Claireville Conservation Area. J.E. was supported by an NSERC postgraduate scholarship. James Fotheringham generously provided stipend support for Y.C.

## Data Accessibility and Benefit-Sharing

Genome assembly and annotation, whole genome sequencing (WGS) data, HiC data and RNA-seq reads were submitted to GenBank (Bioproject: PRJNA1006343). All codes used in analyses have been deposited at https://github.com/YingChen94/Ptriseriata_genome.

## Author Contributions

YC, DRL and SCL conceived of and designed the research. YC and JE conducted fieldwork and acquired the samples. YC, ZS and JE performed the laboratory work. YC and DRL conducted the bioinformatics analyses. YC drafted the manuscript with input from SCL, DRL, VLT and JE. SCL and VLT provided funding and supervision. All authors read, revised and approved the manuscript.

---

**Figure S 1.**
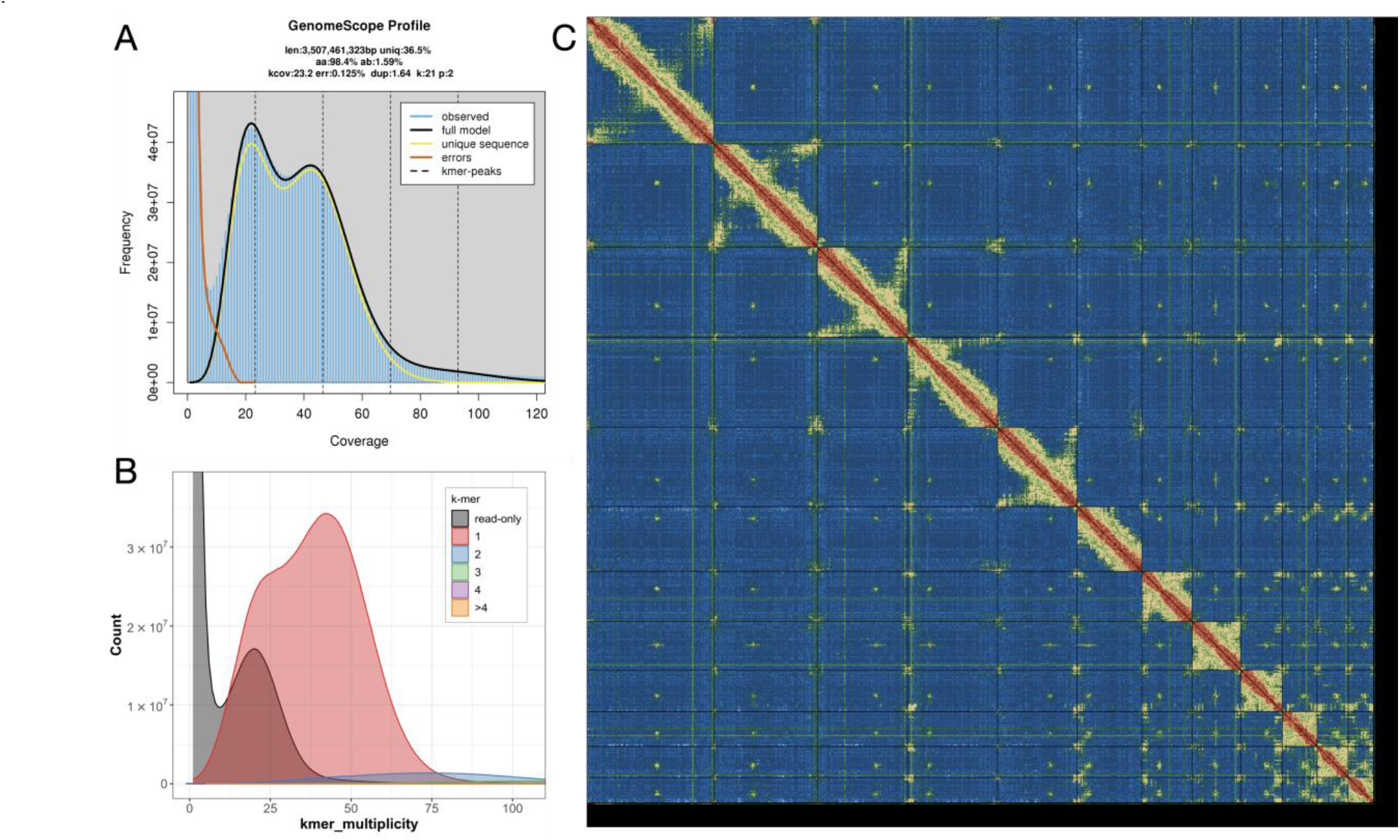
Quality control plots for the western chorus frog (*Pseudacris triseriata*) reference genome assembly. **A.** GenomeScope profile for 21-mers of HiFi long reads. The heterozygote peak is at ∼23x coverage and the homozygote peak is at ∼46x coverage. **B.** Merqury copy number spectra of 21-mers showing haploid and diploid *k*-mers occurred only 1 time in the assembly and thus no haplotypic duplication in the assembly. **C.** Hi-C contact maps showing 12 chromosome-level scaffolds.

**Figure S 2.**
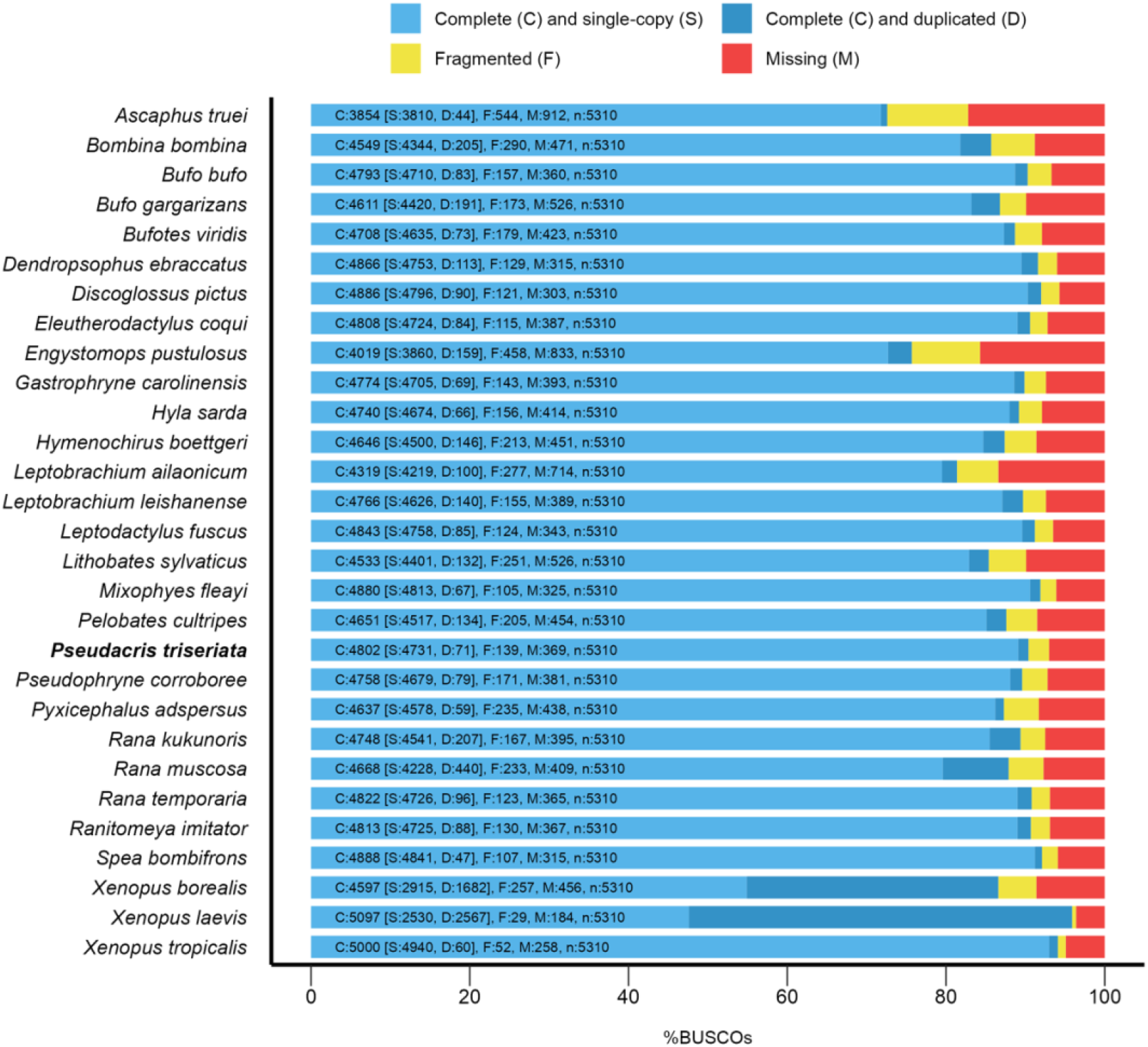
Genome assembly BUSCO assessment results for 29 chromosome-level frog genomes using tetrapoda_odb10 dataset. The western chorus frog assembly from this study is in bold.

**Figure S 3.**
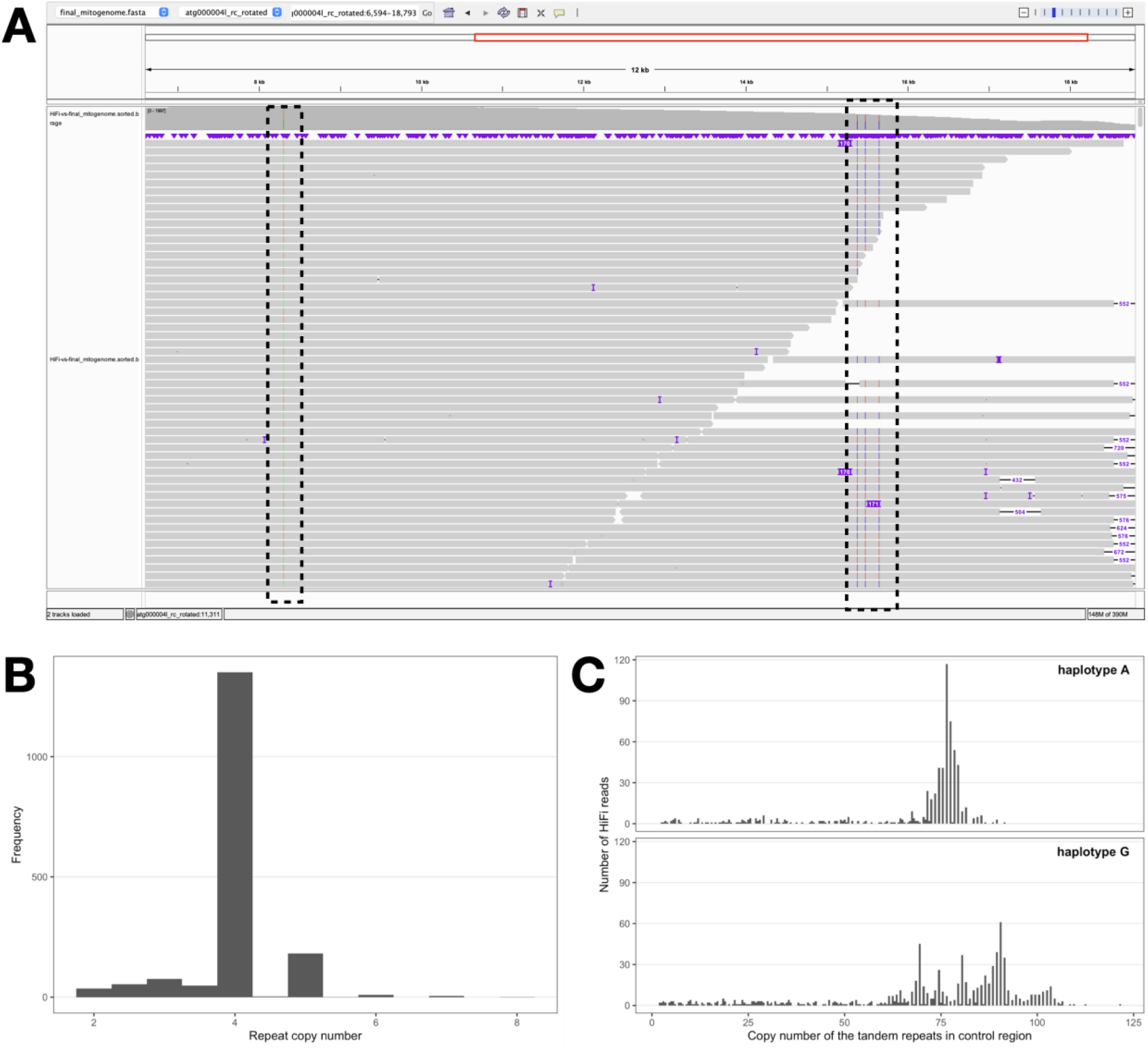
**A.** Multiple haplotypes of the mitochondrial genome are evident in raw HiFi reads visualized in IGV. At position 8302bp (left black box), about 56% reads are G and 44% are A. The right black box shows multiple haplotypes of tandem repeats (repeat unit 170bp) in the control region. **B.** The distribution of copy number for the tandem repeat (repeat unit 170bp) in HiFi reads with 75.7% supporting 4 copies and 10.1% supporting 5 copies. **C.** For two haplotypes (‘A’ vs ‘G’) at position 8302bp, various copy number of the tandem repeats (repeat unit 24bp) are found in raw HiFi reads in the control region.

**Figure S 4.**
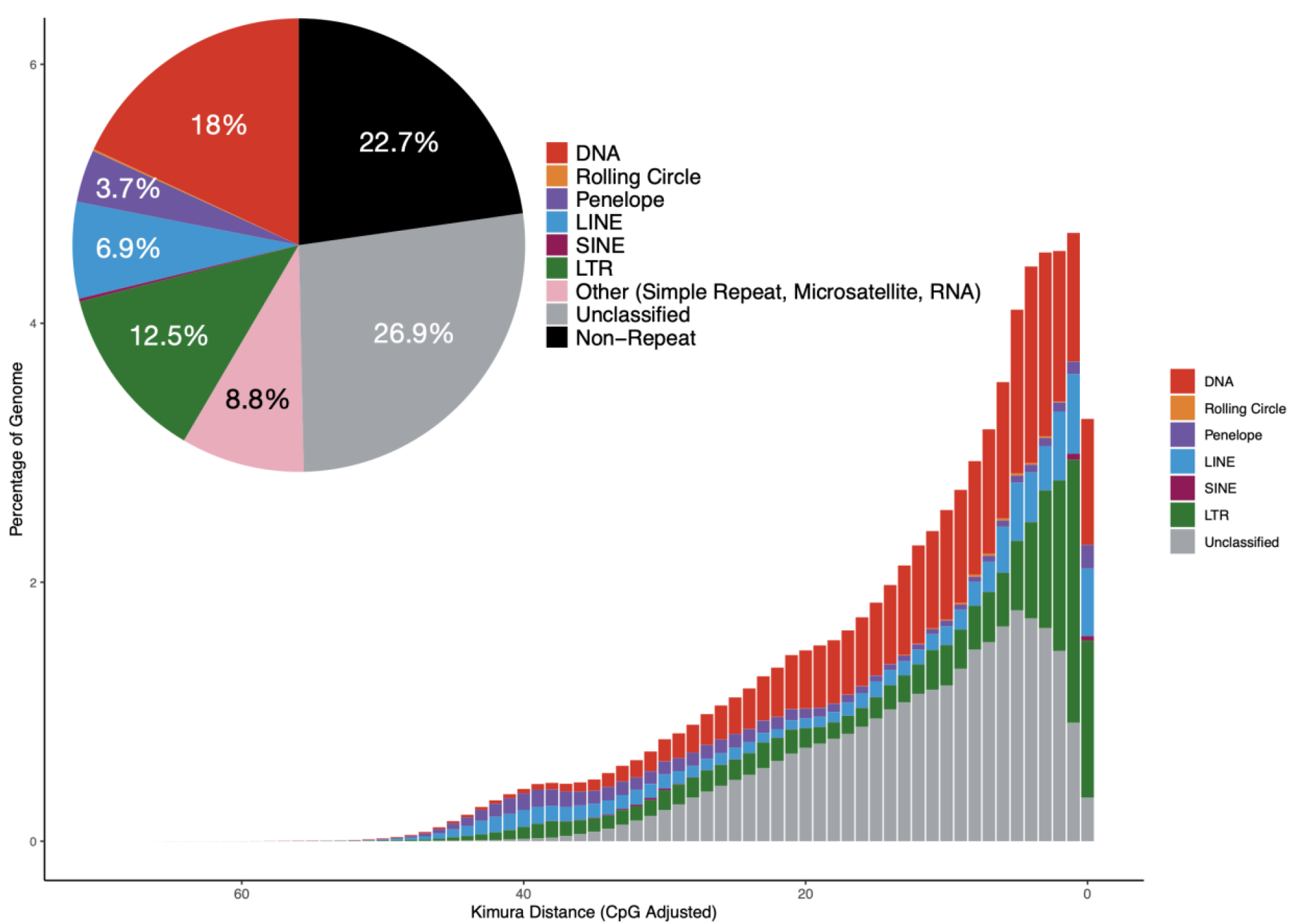
Genome repetitive elements content (pie chart) and the repeat landscape showing the percentage of repeats at different levels of divergence relative to their consensus sequence measured via Kimura distances. The more recent repeat activity (i.e. the repeats with greater similarity to consensus) is closer to the right side.

**Figure S 5.**
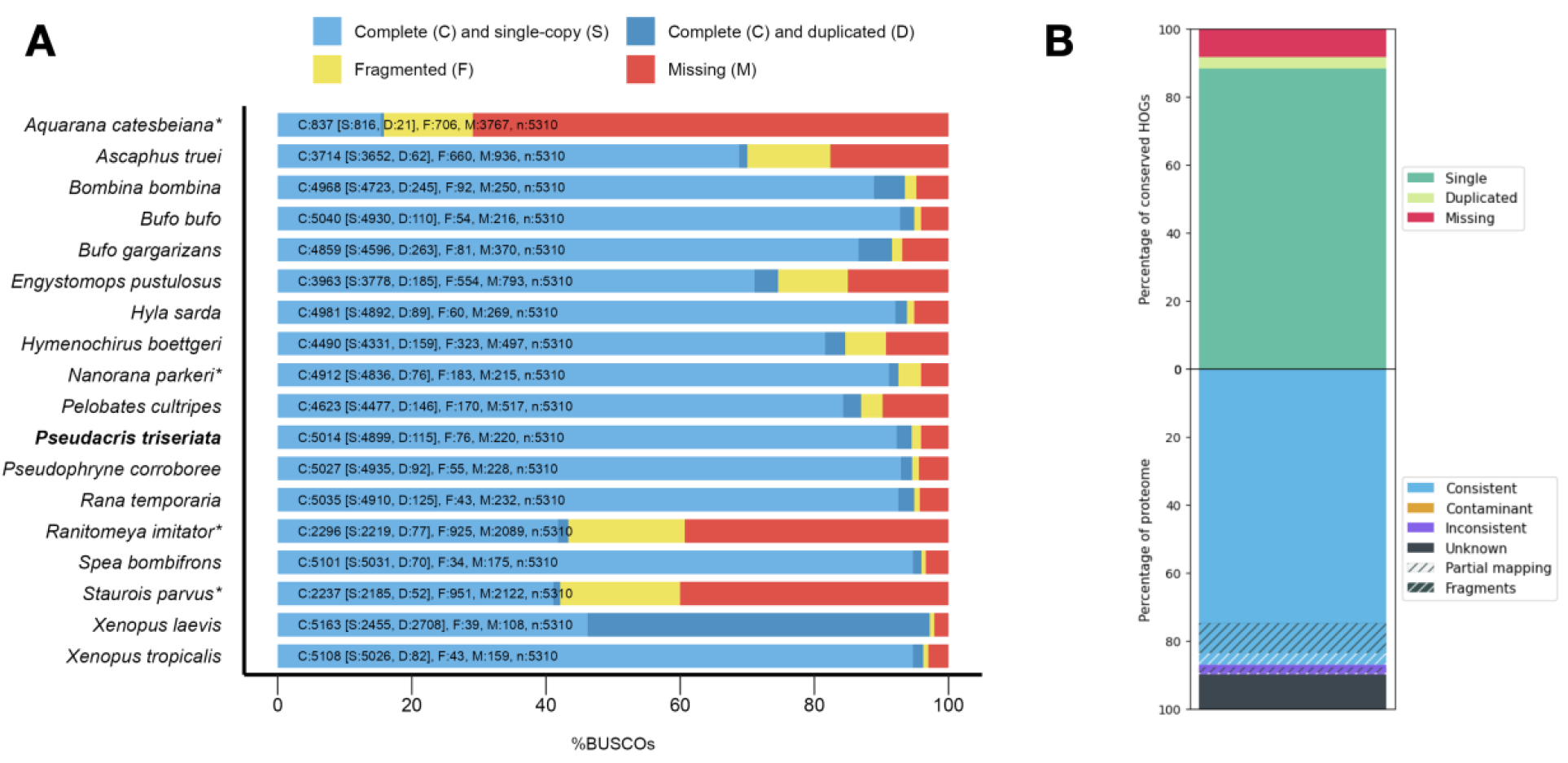
Genome annotation quality assessment. **A.** BUSCO assessment results for the 18 frog genome annotations (only longest isoform per gene was kept in each annotation). Annotations of scaffold-level assemblies are marked with asterisk; the western chorus frog annotation from this study is in bold. **B.** OMArk proteome quality statistics showing high completeness and no evidence of contamination.

